# *In vivo* imaging of individual islets across the mouse pancreas reveals a heterogeneous insulin secretion response to glucose

**DOI:** 10.1101/2020.05.22.111336

**Authors:** Henriette Frikke-Schmidt, Peter Arvan, Randy J Seeley, Corentin Cras-Méneur

## Abstract

While numerous techniques can be used to measure and analyze insulin secretion in isolated islets in culture, assessments of insulin secretion *in vivo* are typically indirect and only semiquantitative. The CpepSfGFP reporter mouse line allows the *in vivo* imaging of insulin secretion from individual islets after a glucose challenge, in live, anesthetized mice, addressing secretion from the pancreas as a whole. Imaging the whole pancreas at high resolution in live mice includes numerous technical challenges. Using rapid-fire imaging in high dynamic range and tiling combined with computer-assisted masked morphometry, we have developed a method to overcome motion blur, and the effects of anesthesia, to be able to monitor and quantify insulin (CpepSfGFP) content simultaneously and longitudinally for the first time in hundreds of individual islets, throughout a glucose challenge.

Through this approach we demonstrate that while isolated islets respond homogeneously to glucose in culture, their response profile differs significantly *in vivo*. Independent of size or location, some islets respond sharply to a glucose stimulation while others barely secrete at all. This platform therefore provides a powerful approach to study the impact of disease, diet, surgery or pharmacological treatments on insulin secretion in the intact pancreas *in vivo.*

## Introduction

With increased insulin resistance, pancreatic β-cells adapt through hypertrophy, hyperplasia and increased insulin secretion. Failure to meet this increased demand leads to Type 2 diabetes (reviewed in 1). Pancreatic exhaustion with depletion of insulin content have been increasingly seen as a leading trigger of islet failure (2-4). While the overall insufficiency of insulin secretion can be measured, little is known about the response of individual islets during normal as well as pathological conditions associated with poor glucose control. Most islet studies are performed by islet purification from the remaining pancreas in order to examine responses *ex vivo* in culture (5). While much can be learned from this approach (6), we know that a wide range of extra-islet signals can have profound effects on insulin secretion. These may include neural signals (7), nutrients (1), hormones (8, 9), and blood flow (10). Thus, isolated islets in culture cannot capture the full complexity that determines insulin secretion *in vivo*. Consequently, it is critical to understand the function of individual islets *in vivo* if we are to assess both the normal ability for glucose to provoke insulin secretion and the islet pathophysiology that is linked to the development of type 2 diabetes.

Several approaches to imaging individual islets *in vivo* have been attempted. Using laparoscopic incisions, investigators have obtained live high-resolution longitudinal imaging of individual islets (10-12). This procedure can be further enhanced through the use of an imaging window to include clusters of islets (13) or using a micro-stage with a stick-type water immersion lens to compensate for the inherent movement due to the breathing of the anesthetized mouse (14).

While each of these *in vivo* approaches has immediate applications for live islet imaging, they are limited by a restricted field of view and the limitations of cellular reporters that can be used to analyze islet function longitudinally, and by the number of islets that can be imaged in each experiment.

Zhu *et al.* recently developed a mouse model with a fluorescent human proinsulin bearing a superfolder GFP-tagged C-peptide (CpepSfGFP) expressed exclusively in β-cells (15). This model allows longitudinal insulin secretion monitoring in live anesthetized mice and originally provided a qualitative appreciation of individual islet response, hinting at differences in glucose response dynamics that could take place between the *in vitro* and *in vivo* conditions (15). We have sought to develop a methodology that allows for *in vivo* imaging and quantification of insulin secretion in individual islets that can capture over time the simultaneous behavior of a much larger number of islets during stimulation. The reporter has been constructed as a transgene that produces only 0.04% of the total insulin made in the islets and does not metabolically alter the animals nor impair the normal functions of β-cells. Nevertheless, the exceptional brightness of the reporter allows for visualization of the accumulated insulin secretory granule fluorescence across the surface of the whole pancreas, within the islets where it is stored prior to secretion. In addition, CpepSfGFP is secreted alongside the normal insulin (16), following the same dynamics. With this approach both expression and secretion of insulin in a number of individual islets can be visualized and quantified in anesthetized mice under various conditions (15).

Described here is a large-scale effort to optimize this approach to make it more reliable, sensitive and quantifiable, tracking the response of large numbers of individual islets over time. We compare the visualization of insulin secretion ex vivo to a wide range of conditions to explore the response of individual islets in situ within the intact pancreas *in vivo*. We find that individual islet response to glucose is quite heterogeneous, implying that a number of factors impact upon glucose response *in vivo*.

## Materials and methods

### Animals

All animal experiments were approved by the Institutional Animal Care & Use Committee at University of Michigan (Animal Use Protocol # PRO00007908). The sfGFP mice were generated as described previously (17) and maintained on a C57BL6/J background. The mice were group-housed and fed standard chow (5LOD irradiated diet, Labdiet, St. Louis, MO, USA). In every round of experiments involving a glucose tolerance test, mice were fasted overnight with food being removed prior to lights out the evening before.

### Islet isolation

Islet isolation was accomplished by collagenase digestion based on the procedures previously described for rat islets (18) with some modifications based on (19). Collagenase was injected into the common bile duct clamped at the ampulla of Vater. After digestion at 37 °C, a series of low-speed centrifugation steps were performed in HBSS-10 % FBS to stop the digestion. The pellet was then filtered through a strainer and centrifuged in Histopaque-1077 (Millipore Sigma, St Louis, MO, USA) under a layer of HBSS. Islets were pipetted from the Histopaque/HBSS interface and hand-picked under an inverted microscope.

### Cell culture and glucose-stimulated insulin secretion *in vitro*

After isolation, the islets were kept four days in culture at 37 °C with 95% humidity in media (RPMI-1640 with L-Glutamate, 10% FBS, Penicillin/Streptomycin, Minimum Essential Medium (MEM) non-essential amino acids, HEPES (Gibco, Waltham, Massachusetts, USA]). For imaging, islets from four mice were pooled and redistributed in four different wells (50 islets per well). The islets were embedded within a thin layer of 500 μL of Rat Type I collagen (Corning, Bedford, MA, USA), as previously described (20, 21). After polymerization, the gel was overlaid with 500 μL of culture media and allowed to rest for an additional day before stimulation and imaging.

### Islet glucose stimulation and imaging

Prior to glucose stimulation, the media for the islets in collagen was replaced with HEPES-buffered, Kreb-Ringer solution with 1 mM glucose for 3 hours, replacing the media after 1.5 hours. The plates for the islets were placed on a mat heated at 37 °C on a H101A ProScan motorized stage (Prior Scientific Instruments Ltd. Cambridge, United Kingdom) under a Nikon AZ100 multizoom microscope (Nikon Instruments Inc., Melville, NY, USA). The wells with the CpepSfGFP islets in collagen were then imaged before stimulation and then every 5 minutes throughout the experiment with a CoolSNAP (Teledyne Photometrics, Tucson, AZ, USA) 12-bit cooled CCD camera, through the NIS-Elements AR software (version 5.0.2; Nikon Instruments Inc. Melville, NY, USA). After 15 minutes of imaging in low glucose, the medium was then replaced with 500 μL of HEPES-buffered, Kreb-Ringer solution with 52 mM glucose to bring the final concentration of glucose (media+gel) to 26 mM. After 15 minutes, the medium was replaced again with 1 mM glucose HEPES-buffered, Kreb-Ringer solution for 15 minutes, and then, finally with HEPES-buffered, Kreb-Ringer solution with 60 mM KCl (to bring the final concentration of media+gel to 30 mM KCl). At the end of each step, media were collected, stored at −80 °C, and insulin measured by ELISA.

### Islet imaging analysis

The high dynamic range 12-bit images from the experiment were analyzed Fiji (version 2.0.0-rc-69/1.52p), a distribution of ImageJ (22-24). As illustrated in Figure 2, each image was duplicated. One of the images underwent background subtraction using the rolling ball method, and thresholding to create a mask. The measurement settings were adjusted to redirect the collection of the integrated density of the areas thresholded on the mask on the original unaltered copy of the image. “Particle” analysis in Fiji allowed the quantification of the integrated density (sum of the intensity of all the pixels in the area delimited by the mask) the of individual islets or small groups of islets, displaying the corresponding label on each image (see flowchart in Figure 2). The same process was repeated for each image of individual wells throughout the experiment. All the data collected could then be parsed in a FileMaker database (version 17, Claris International Inc. Santa Clara, CA, USA). Using the labels displayed in the image, the islets were visually mapped from image to image, allowing to create a relational database of all the data (Supplemental Figure 1) and a trace of the individual islet intensity over time. Through the relational database, the data output could then be simply tabulated and graphed, providing a trace of the integrated density of each individual islet over time, as illustrated in Figure 3.

**Figure 1:**
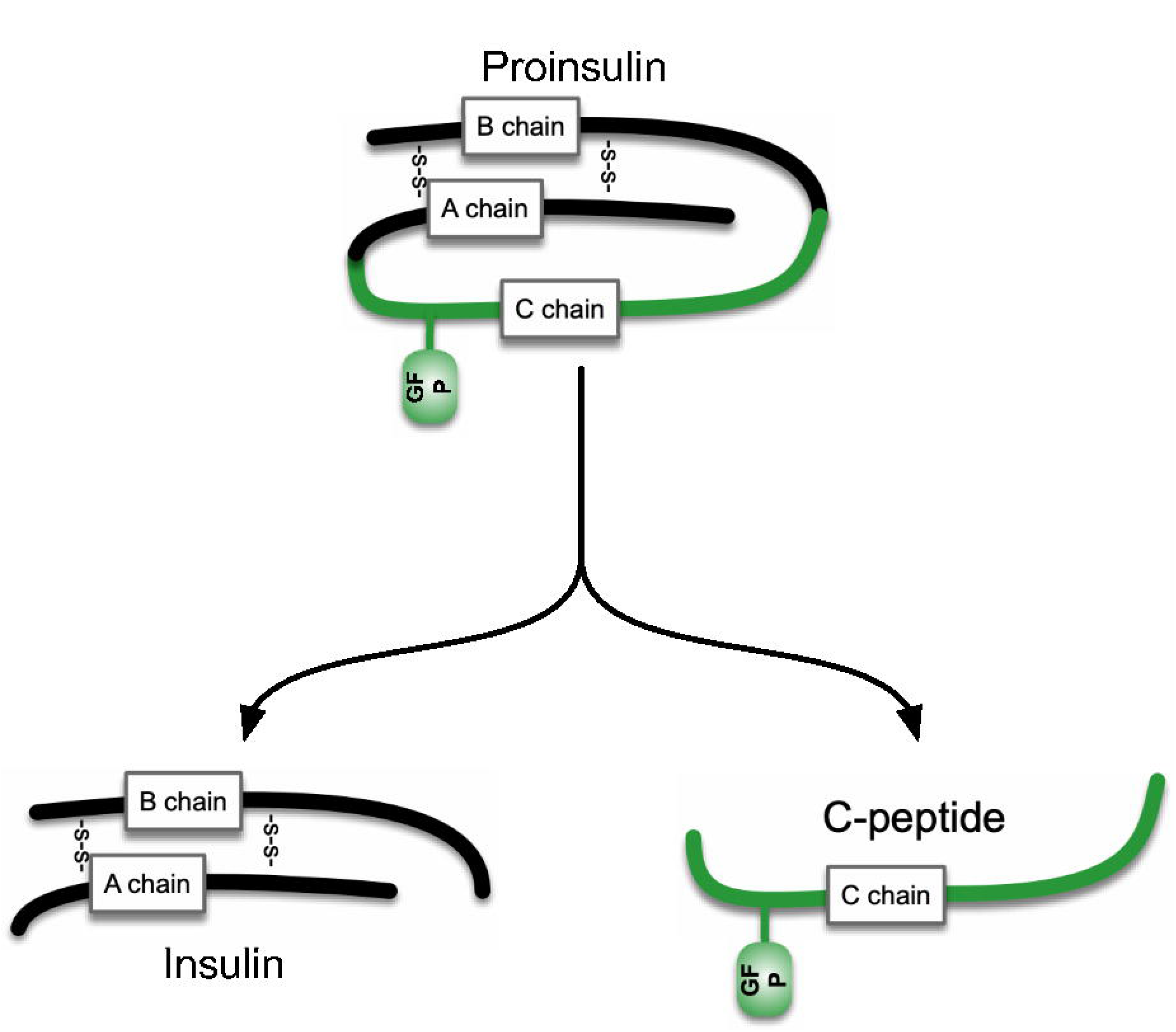
CpepSfGFP diagram. Diagram of the structure of the proinsulin with the super folder GFP (sfGFP) tag attached to it. Processing the proinsulin separates the normal insulin (left) from the CPeptide with the sfGFP tag attached to it (right).

**Figure 2:**
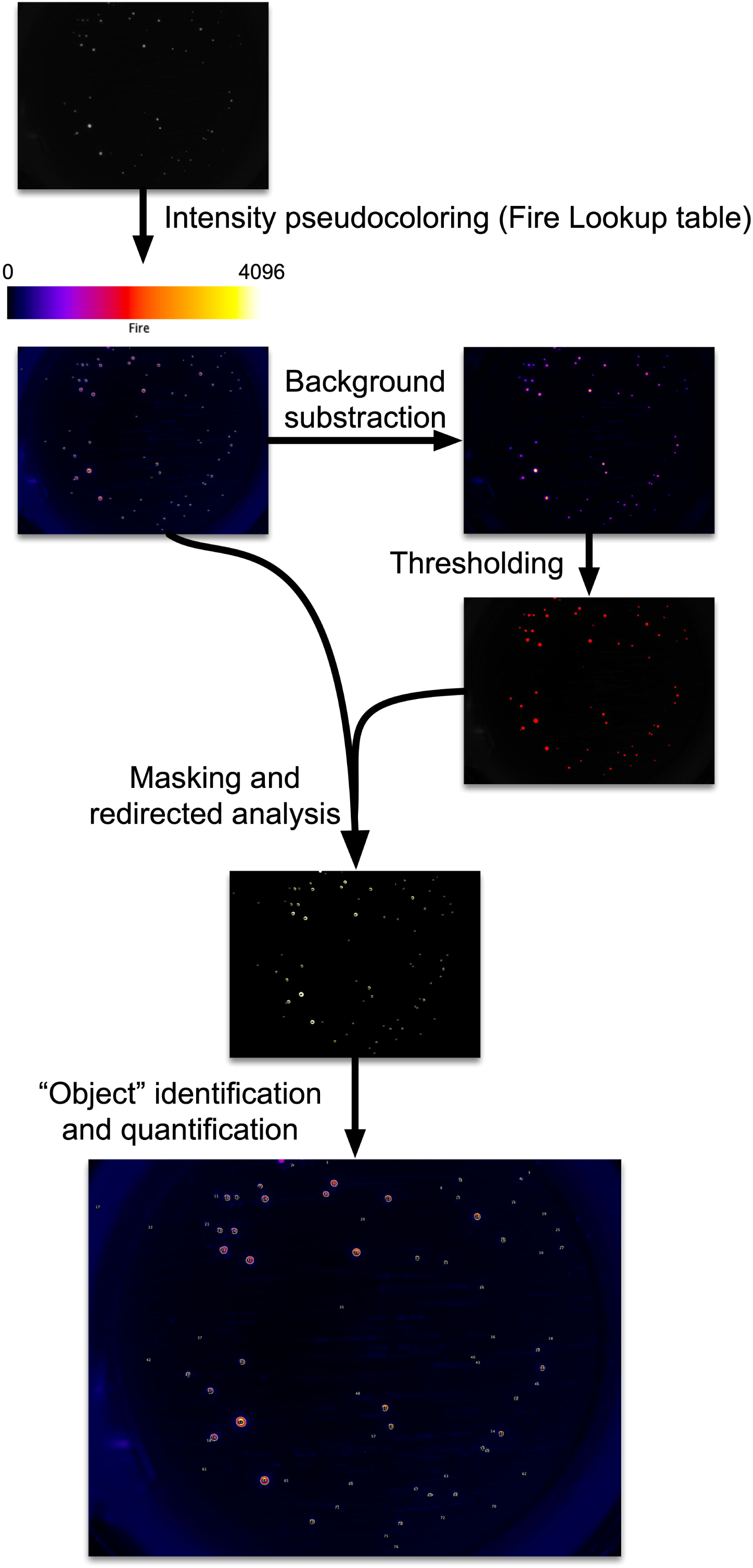
CpepSfGFP imaging on islets in culture. Flowchart of the imaging and analysis of the CpepSfGFP islets in culture. Captured images undergo pseudo-coloring using the “Fire” lookup table (scale on top of the image). Background subtraction and thresholding are used to generate a mask to be used for islet identification on the original image. Each islet or small cluster of islets (defined as “objects”) can be quantified and compared over the different time points.

**Figure 3:**
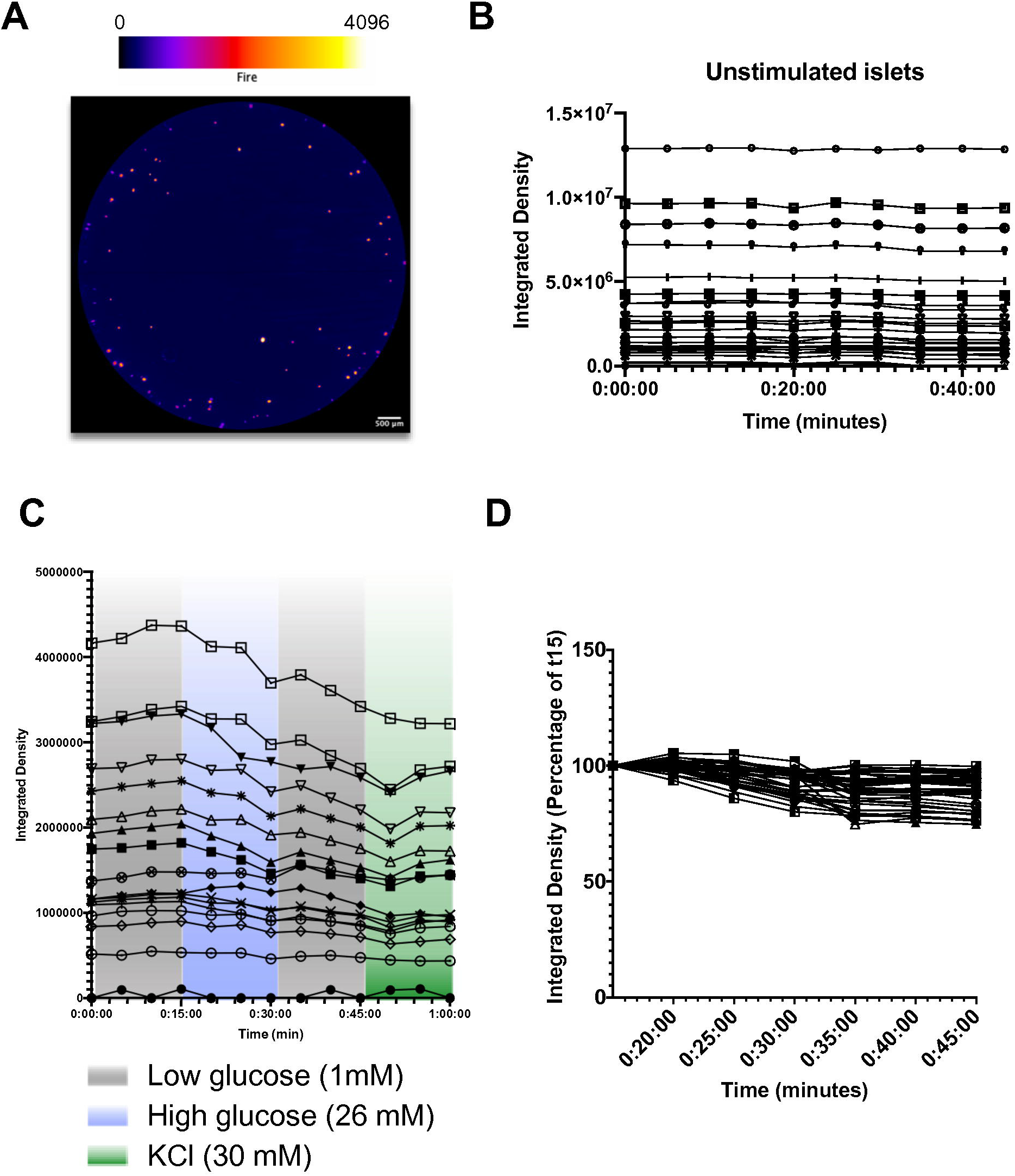
CpepSfGFP profiles in islets in culture. **A.** pepSfGFP islets in culture in collagen with intensity-based “Fire” pseudocoloring (scale on top). **B.** Integrated density of the unstimulated individual CpepSfGFP islets over 45 minutes of culture. **C.** Integrated density of CpepSfGFP individual islets in culture during a GSIS in low (1 mM) glucose, then high (26 mM) glucose, low glucose again and finally KCl (30 mM). **D.** individual islets from Figure C represented as a percentage change from t15 (time at which the islets are stimulated with glucose). Scale bar = 500 μm

### Insulin measurements

Insulin was measured using the Ultra Sensitive Mouse Insulin ELISA Kit (Crystal Chem, Elk Grove Village, IL) according to the manufacturer specifications.

### Analgesia and anesthesia

Ostilox was used for analgesia at a dose of 0.1 mg.kg^-1^ administered two hours prior to the individual procedure. Anesthesia was either isoflurane or pentobarbital (Euthasol Virbac AH, Inc., Fort Worth, TX) given in an induction dose of 81 mg.kg^-1^ followed by a maintenance dose of 20 mg.kg-1 given every 20 minutes until the end of the procedure. Acetaminophen (Sigma Aldrich, St. Louis, MO) was administered at a dose of 0.1 mg.kg-1 for gastric emptying measurements.

### Surgical procedures – sandwiching the pancreas

After being anesthetized, the mouse was placed on a heating pad maintained at 37 °C. The left side of the mouse was shaved and placed in between 2 hollow copper tubes of. 1.5 cm of diameter. Copper conducts heat efficiently, and the tubes prevent heat loss from the sides of the mouse. They also served as a heated platform supporting the microscopy slides above the animals. An incision was made just above the spleen but below the level of the diaphragm. The pancreas was gently exteriorized and placed on the warmed microscopy slide resting on the copper tubes. It was ensured that the pancreas was flat and in a single layer across the slide. Plastic spacers 3 mm thick were placed on each side of the pancreas. Hypromellose ophthalmic demulcent solution (2.5%) (Goniovisc; HUB Pharmaceuticals, Plymouth, MI, USA) warmed at 37 °C was applied to the surface of the pancreas before a coverslip was placed over the spacers, “sandwiching” the pancreas at uniform thickness between the slide and coverslip (see diagram in Figure 4). The gel prevents tissue drying while offering optimized light conductance for the sfGFP as previously illustrated (25).

**Figure 4:**
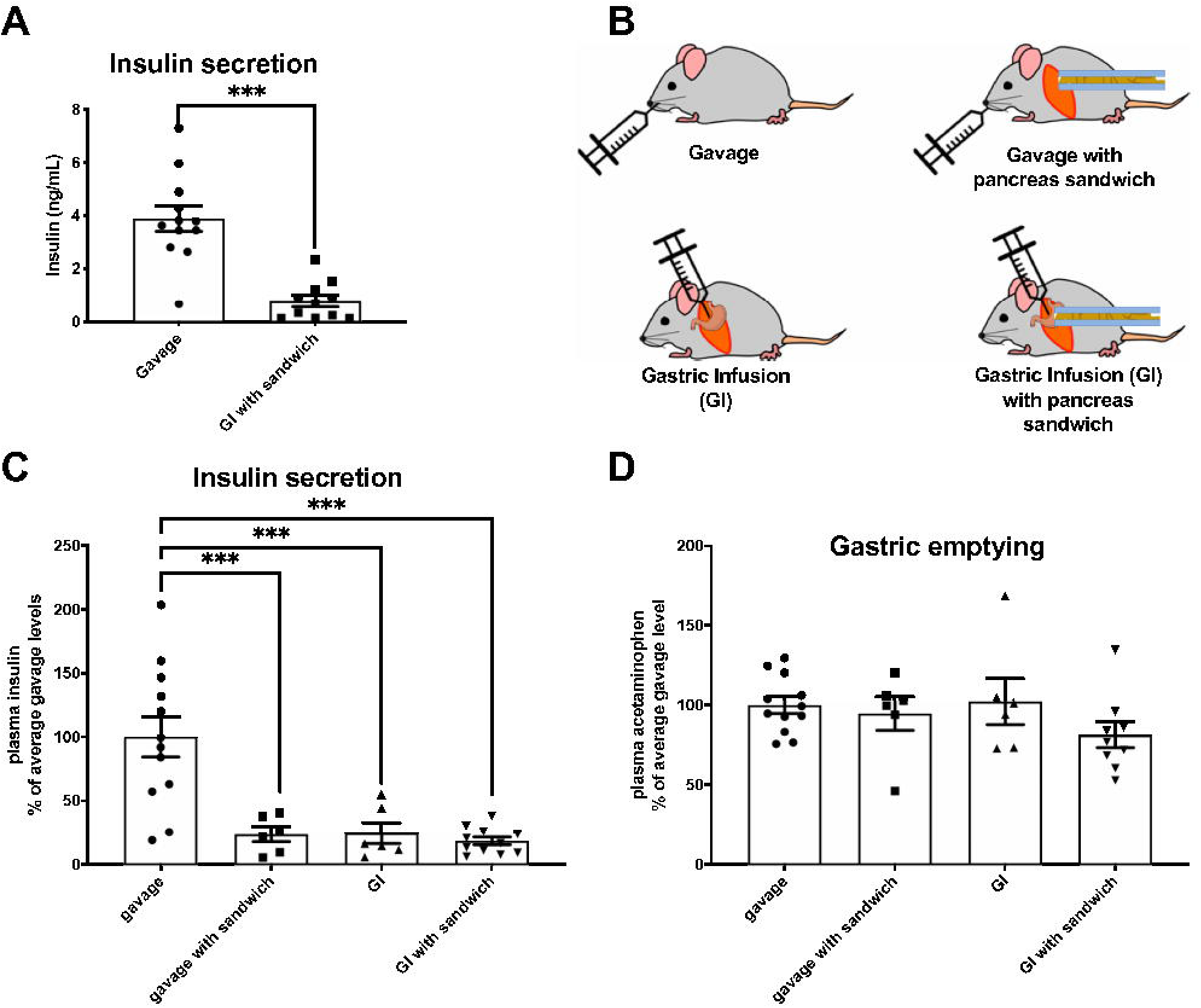
Impact of surgery. **A.** Glucose-stimulated insulin secretion on mice stimulated by gavage or by gastric infusion after having the pancreas exteriorized. **B.** Diagram of the different conditions used for analysis in panels C and D. Insulin secretions (**C**) and gastric emptying (**D**) in mice that were glucose-stimulated by either gavage, gavage with pancreas exteriorization (sandwiching), gastric infusion and gastric infusion with pancreas exteriorization (sandwitching). *** p <0.001

**Figure 5:**
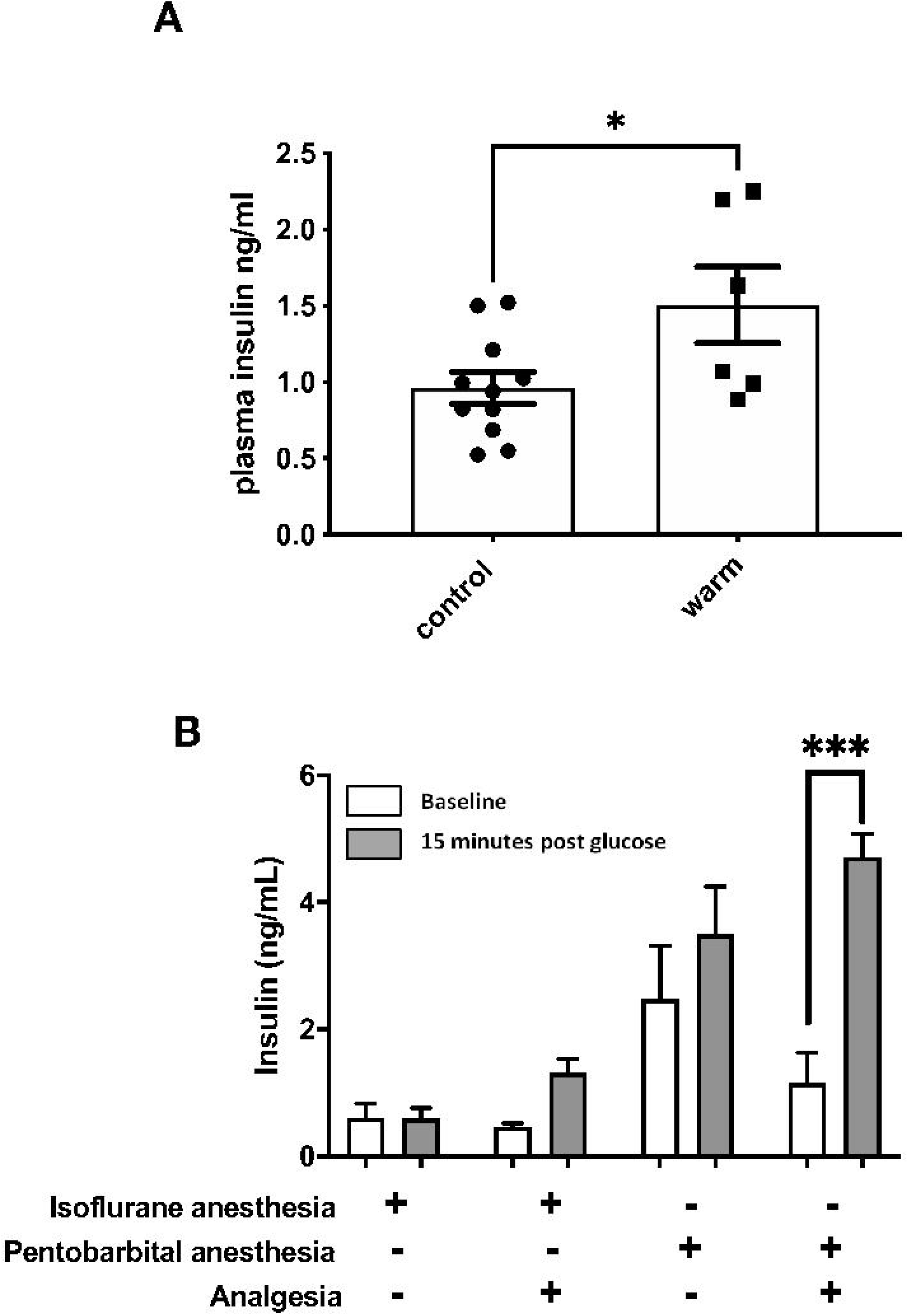
Impact of temperature, anesthesia and analgesia. **A**. GSIS in anesthetized mice with the pancreas exteriorized and sandwiched on a slide on plastic spacers (control), or on copper tubing (warm). **B**. Comparison of GSIS on animals with either isoflurane or pentobarbital and with or without analgesia. * p <0.05, *** p <0.001

The heating pad with the mouse was then placed on top of the H101A ProScan motorized stage (Prior Scientific Instruments Ltd. Cambridge, United Kingdom) under the Nikon AZ100 multizoom microscope (Nikon Instruments Inc., Melville, NY, USA) for imaging. Tail vein blood was obtained for glucose measurements and plasma insulin.

### Whole pancreas live imaging

The pancreas was subsequently also imaged on the AZ100 microscope using a CoolSNAP (Teledyne Photometrics, Tucson, AZ, USA) 12-bit cooled CCD camera, through the NIS-Elements AR software (version 5.0.2; Nikon Instruments Inc. Melville, NY, USA). The objective (5x) and secondary magnification (1x) were chosen to maximize the depth of field and fluorescence transmission. At this magnification, the imaging of the entire pancreas required tiling. To increase the chances of capturing without motion blur, NIS-Elements AR was programmed to capture 5 images per field in 12 bit, 250 ms apart before moving to the next field. Tiling was acquired with a 15% overlap.

After the baseline imaging, a flat dose of 75 mg glucose was administered either by gavage or intra-gastric injection through a catheter placed distally from the incision. Every 5 minutes the pancreas was imaged followed by blood glucose measurements. The procedure ended at 15 minutes after glucose administration, at which time blood was obtained for plasma insulin measurement as well. The mouse was euthanized by cervical dislocation.

### CpepSfGFP image analysis and islet quantification

The images were extracted from the NIS container using NIS Viewer 4.11.0 (Nikon Instruments Inc. Melville, NY, USA) and then reviewed in Fiji individually to select only one image without motion blur per field (see flowchart in Figure 6). All the images of a mosaic were then reassembled in Fiji using the Grid/Collection stitching plugin from Stephan Preibisch, freely available on the ImageJ repositories (https://imagej.net/Image_Stitching). Similar to what was described above for *in vitro* islet imaging, the image underwent the same analysis process: duplication, background subtraction, thresholding, redirection of the particle analysis on the original unaltered image to quantify individual integrated densities and labelling before collecting and relating all data in a FileMaker database to output the integrated density of each individual islet (or cluster of islets) over time throughout the glucose challenge. Any suspicion of technical artifacts (close proximity to the edge of the slide, imaging issues, proximity of a bubble in the optical gel, etc.) led to the elimination of the islet from further analysis.

**Figure 6:**
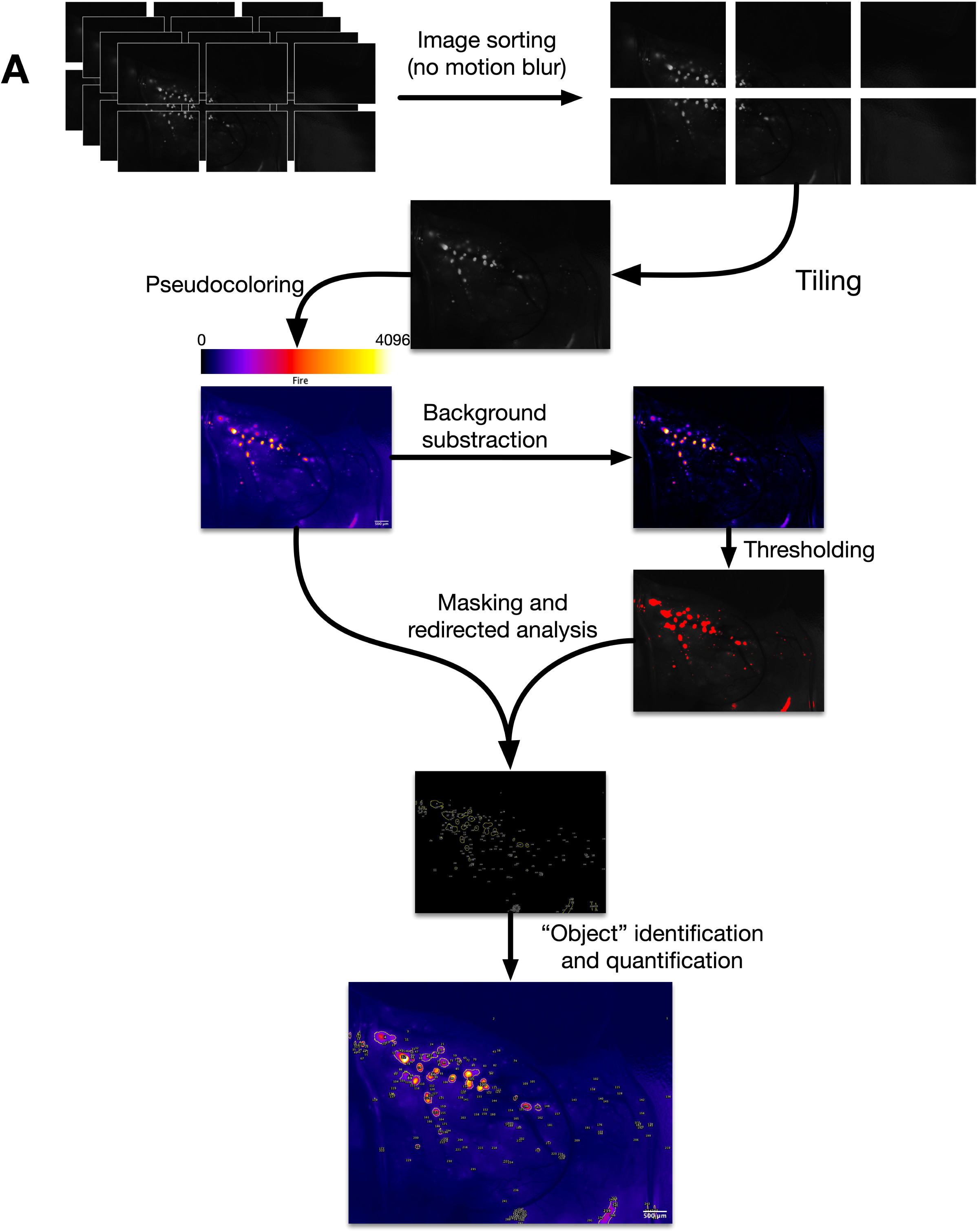
Live CpepSfGFP islet imaging. Flowchart of the imaging and analysis of the CpepSfGFP islets *in vivo* in anesthetized mice. Rapid-fire imaging allows the capture of multiple images in each position. The images are then filtered to eliminate the ones with visible motion blur and tiled to recompose a complete high resolution view in 12 bit of the entire pancreas. The images undergo pseudo-coloring using the “Fire” lookup table (reference scale on top of the image). Background subtraction and thresholding are used to generate a mask to be used for islet identification on the original image. Each islet or small cluster of islets (defined as “objects”) can be quantified and compared over the different time points. Scale bar = 500 μm

### Statistical analysis

Comparisons that passed a normality test were assessed through paired or unpaired t-tests. If normality tests failed, comparisons were analyzed using a Mann-Whitney test for unpaired data. For multiple comparisons, we used either 1-way ANOVA with Tuckey’s multiple comparison test, or 2-way ANOVA following the Holm-Šídák method.

## Results

### CpepSfGFP islets respond synchronously to a glucose challenge *in vitro*

The CpepSfGFP reporter is a bright reporter that reflects insulin content. When stimulated with glucose, the islets secrete insulin/CpepSfGFP and the integrated density of the reporter fluorescence goes down. In their original report of the CpepSfGFP model, Whu *et al.* were unable to detect intensity changes in isolated islets free floating in culture and with 8-bit imaging upon glucose stimulation (15).

In order to be able to track individual islets in culture over the duration of a glucose challenge, islets were immobilized in culture in a collagen gel similarly to what has been previously used for embryonic pancreatic rudiments in culture (20, 21, 26-28). The collagen is considered to be mostly inert and doesn’t affect insulin secretion. Its wide pores allow for nutrient and glucose rapid diffusion to the embedded tissue.

Islets were embedded in collagen with media added on top of the gel layer after polymerization. As illustrated on Figure 2, fluorescence in the wells from individual CpepSfGFP islets was easily detected. Using a high dynamic range camera (12-bit imaging), both low and high intensity islets could be imaged and quantified without clipping of the signal (represented by pseudo-color with an intensity-based lookup table in Figure 2). Islets could then be imaged every 5 minutes over the course of GSIS. Masking after background subtraction helped to delineate individual islets or small clusters of islets. Using these masks, the intensity of the signal on the original images derived from insulin/CpepSfGFP content in the individual islets over time were quantified throughout the experiment (represented in Figure 2). In the experiments presented in Figure 2 and 3, the density of islets in the well was chosen to minimize the risk of having islets aggregated together, and with our mask, all images allowed clear delineation of individual islets.

Islets kept in culture in low glucose (1 mM) did not display any change of intensity over time (quantified over time in Figure 3B). Regardless of their size or location, their levels remained stable, illustrating that the imaging conditions did not lead to any appreciable photo-bleaching. Additionally, when challenged with high glucose (26 mM) and thereafter with KCl (30 mM), essentially all islets responded to these stimuli as measure by a loss of SfGFP signal intensity from the islets (Figure 3C). After a 15 min challenge with high glucose, average islet SfGFP signal intensity declined ∼10% from initial values, and after a subsequent 15 min stimulation by depolarization with KCl, the average SfGFP signal intensity declined to ∼12.5% from starting values. Importantly, the response of the islets appeared independent of islet size, consistent with the notion that all islets in culture secrete a small fraction of their available insulin into the media in response to stimulation with high concentrations of glucose or potassium-mediated depolarization (Figure 3D).

### Surgical distress blunts insulin response *in vivo*

Visualizing the CpepSfGFP islets in live mice requires a laparotomy and exteriorization of the pancreas. Any form of surgery, even under anesthesia, triggers a stress response with an increased secretion of pituitary hormones and activation of the sympathetic nervous system (29). In a preliminary set of experiments, the insulin secretion of animals that underwent laparotomy for pancreas imaging were compared to control mice that had been gavaged. Figure 4A demonstrates that while animals that were anesthetized and had surgery for pancreas imaging still exhibited glucose-stimulated insulin secretion, the magnitude of the response was fractionally smaller than that of the reference group (non-anesthetized mice subjected to glucose stimulation through gavage).

To avoid gavaging of anesthetized mice (which poses technical challenges), we tested the use of gastric infusion by inserting a catheter into the stomach as part of the surgical procedure, prior to pancreas externalization.

In order to distinguish the effects of the surgery itself from the method of glucose stimulation, the animals were split in four groups 1) gavaged mice, 2) gavaged mice with pancreatic exteriorization, 3) mice that underwent a simple laparotomy without pancreatic exteriorization and had a glucose stimulus through gastric infusion and finally 4) mice that underwent laparotomy, pancreatic exteriorization and glucose stimulation by gastric infusion (see diagram in Figure 4B). In all cases, regardless of the method used for glucose stimulation, all three groups that had a laparotomy had a blunted insulin secretion in response to glucose (Figure 4C). In addition, all four groups had near identical gastric emptying rates (Figure 4D).

### Temperature control, anesthesia and analgesia improve the magnitude of insulin secretion in response to glucose

While the mice were always kept on a 37 °C heating pad during the surgical-imaging procedure, exteriorization makes the pancreas itself subject to additional heat loss. Despite pre-warming the slides to 37 °C for imaging, we noted that the temperature on the slide dropped to 22 °C within a few minutes. In order to test the possibility that this drop in temperature could blunt insulin secretion, we replaced the plastic spacers defining the pancreatic sandwich with copper tubing. Copper has a thermal conductivity of 399 W.m^-1^.K^-1^ (over 400 times better than most plastics). The tubing was placed alongside the back and the abdomen of the mouse, reinforcing core temperature control on the mouse itself, and increasing the temperature on the slide to 36.8 °C. We therefore compared insulin secretion with the plastic spacers or with copper tubing and noted a ∼1.5-fold increase in insulin secretion (Figure 5A).

Isoflurane is one of the most widely used anesthetics in mice. Induction and recovery are fast, and the risk of accidental death is low. Although the effects of anesthesia on insulin secretion have been previously studied (30-34), most anesthetic agents can also blunt insulin secretion to glucose to some degree (30, 31, 34). Nevertheless, pentobarbital has been reported to have less impact on glucose tolerance tests and insulin secretion (34). We therefore compared anesthetic agents and analgesia (using copper tubing for temperature control) to further examine the impact of the laparotomy on glucose-stimulated insulin secretion *in vivo*. Figure 5B demonstrates that pentobarbital can in part restore proper insulin secretion compared to isoflurane, and that analgesia further ameliorates insulin secretions in response to a glucose challenge. Together, temperature control, analgesia and the use of pentobarbital for anesthesia allowed recovery of stimulated insulin secretion from the exteriorized pancreas that best resembles what is typically observed from non-anesthetized mice.

### Optimized platform for CpepSfGFP GSIS imaging in anesthetized mice

Imaging a live pancreas under fluorescence in an anesthetized mouse presents additional challenges compared to imaging islet CpepSfGFP secretion *in vitro*. The live animals are breathing so that the abdomen is constantly moving under the camera. This can lead to significant motion blur, complicated by the fact that the large pancreas area cannot be imaged in only one field without a dramatic loss of magnification/resolution. Indeed, in order to be able to simultaneously track as many islets as possible, the images need to be acquired with a wide depth of field, keeping as many islets as possible in focus.

To cover the entire area of the tissue, a microscope and objective was chosen with a long working distance and wide depth of field, allowing image capture of the entire pancreas on a motorized stage by tiling ∼6 fields, while maintaining most imaged islets within the focus range. To minimize motion blur, we programmed the microscope to take 4 images in rapid-fire from each field chosen by the motorized stage. Only one of the four images was subsequently chosen to maintain focus and synchronize the image capture with the breathing cycle of the animal. Tiling allowed us to recompose a complete overview of the tissue (schematized in Figure 6). The remainder of the imaging analysis used the same approach as that for isolated islets *in vitro* in collagen. Using this platform, we were able to track over the glucose challenge up to 200 individual islets, and small clusters of islets.

### CpepSfGFP islets respond heterogeneously to a glucose challenge in live, anesthetized mice

While isolated islets respond in a coordinated fashion *in vitro*, we were surprised to find that islet response to intragastric glucose is vastly heterogeneous *in vivo*: some islets dropped dramatically in intensity while others did not seem to respond at all (Figure 7A). Looking at small groups of islets at a higher magnification suggested no consistent topographical impact in determining islet response to high glucose stimulation. In small groups of islets, we often saw some islets responding sharply while immediately adjacent islets exhibited little change in intensity (Figure 7B). A representation of the islet dynamics in terms of the proportion (percentage of the changes based on t0) illustrates the wide range of response observed *in vivo* (Figure 3C, D). Plotting the islet response in a logarithmic scale based on integrated density change ratio (on the vertical axis, in log scale) and area (on the horizontal axis) demonstrated no discernable relationship between islet size and insulin secretion, ruling out a size effect (Figure 7D). A few of the small islets seemed to show actual increase in insulin content, which could relate to actual insulin biosynthesis or the fact that smaller islets are more susceptible to measurement variability inherent to the movement of the pancreas in three dimensions. As illustrated in Figure 3D, stimulated islets in culture respond quite homogeneously with a very tight SEM (Figure 7F). *In vivo*, the islet responses are vastly more heterogeneous, with a much wider SEM (Figure 7F). Comparing the loss (of integrated density) of islet CpepSfGFP upon high glucose stimulation *in vivo*, the overall response of all islets together averaged ∼8% of total insulin/CpepSfGFP content, which was quite similar to that seen *in vitro* (Figure 7E). However, the comparison of the SEM of the responses of individual islets in culture and *in vivo* clearly reflected a greater heterogeneity under the latter condition (Figure 7F).

**Figure 7:**
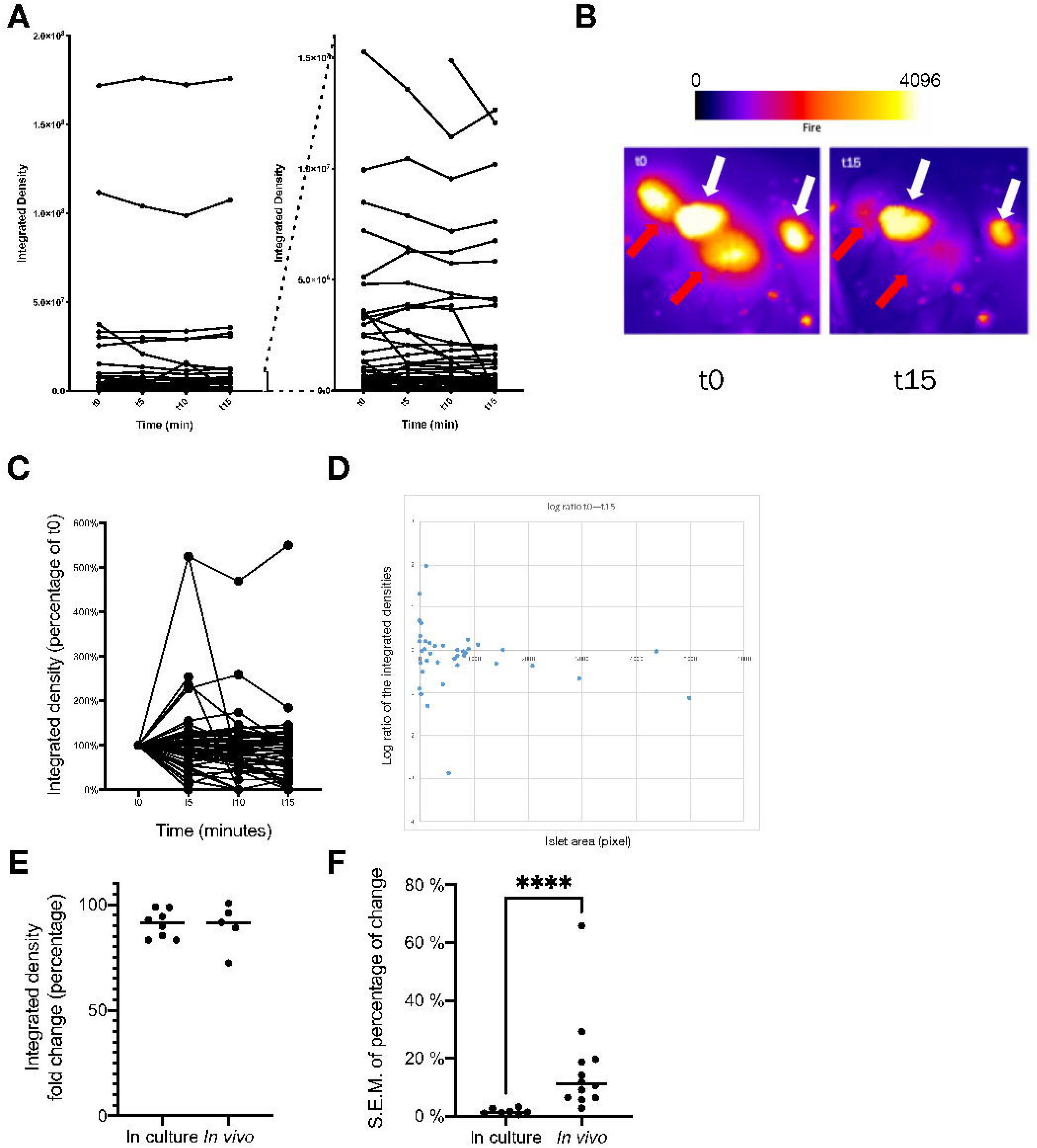
CpepSfGFP profiles on islets *in vivo* in anesthetized mice. Expression profiles of the CpepSfGFP islets *in vivo* in anesthetized mice after a glucose stimulation (gastric infusion) at t0. **A**. expression profile of all individual islets (left) or a subset of smaller islets (different scales on the right) over 15 minutes. **B**. Pseudo-colored islets (Fire lookup table, scale above) before stimulation (t0) and after 15 minutes (t15). The white arrows indicate islets that exhibit only modest changes in 15 minutes while the red arrows highlight adjacent islets that change dramatically over the same time course. **C**. Integrated density of the individual islets expressed as a percentage at t15 relative to their value at t0. **D**. Log ratio representation of the islet response based on the integrated density on the vertical axis and their area (expressed in pixel) on the horizontal axis. **E.** Comparison of the total islet response to the glucose challenge in culture and *in vivo* (percentage change of the sum of all integrated densities in a culture well or a pancreas). **F**. SEM of the islet response (comparing the distribution of individual islets response expressed as a percentage at t0 compared to t15) of islets in culture compared to islets imaged *in vivo*. **** p <0.0001

## Discussion

The development of type 2 diabetes results from reductions either in islet mass, function, or both. Dramatic changes in islet mass can affect the long-term risk of developing diabetes. Pancreatic morphometry after immunostaining has been a long-established gold standard to assess β-cell mass. On the other hand, very few functional tests have been available for *in vivo* studies, in particular when attempting to address islet response in the whole pancreas. In 1987, Stefan *et al.* sampled the pancreas by immunostaining after a glucose challenge and hypothesized that insulin secretion was heterogeneous within an islet, as well as between islets, depending upon where they were located (35). Their approach was purely qualitative and provided limited sampling, but it was a first attempt to address potential heterogeneity of insulin secretory responses of islets *in vivo*.

In 2016, Zhu *et al.* described a mouse model that could allow monitoring of insulin storage and secretion *in vivo* and *in vitro* (15). The original method allowed for the crude analysis of the dynamic of secretion for groups of islets at the time as well as important qualitative aspects. However, we discovered, and devised means to circumvent, multiple technical obstacles that affect insulin secretory responsiveness *in vivo*, and the ability to accurately image and quantify a wide range of the insulin content within multiple islets across the surface of the pancreas over a time course. Here, we have presented a means to significantly improve the original platform to longitudinally monitor hundreds of islets at a time *in vivo* in anesthetized mice. Using this paltform, we have been able to visualize for the first time the differences in the response of islets *in vivo* compared to what had been described using *in vitro* approaches. While islets *in vitro* more consistently secrete a small fraction of their individual insulin content in response to high glucose, islets *in vivo* display drastically different responses throughout the pancreas (Figure 7).

Investigators have previously attempted to image secretion from live individual islets (12), or groups of islets (12) in a number of limited and indirect ways. In this manuscript, we have used a fluorescence reporter that allowed us to directly monitor insulin/CpepSfGFP secretion in hundreds of individual islets at a time, repeatedly, throughout a glucose challenge.

We note several caveats to our studies. Exteriorizing the pancreas allows *in vivo* imaging for most of the body and tail of the pancreas but imaging the head of the pancreas was not physically possible without significant mechanical stress on the duodenum and stomach. In addition, smaller islets that are embedded deep within the pancreas are less likely to be imaged. In order to minimize this problem, we used an objective with a wide depth of field, maximizing the number of islets that are within focal range. The SfGFP reporter is extremely bright and allows capture of islets that could be deeper from the pancreatic surface, with the caveat that the thickness of overlying non-islet tissue above can weaken fluorescence signal, diffuse the emitted light, and thus blur the shape of the islets. In addition, the mice are anesthetized and breathing, leading to movements of the pancreas. Having the pancreas resting on a heated slide somewhat limits the magnitude of movements of the tissue, but it could still affect imaging of individual islets. While large islets are mostly unaffected by these small movements in 3D, the effect is not always negligible on smaller islets. These caveats do not affect the overall integrated density of the captured images, although they can affect the precise estimation of islet size and the degree of intensity changes on the smallest islets, which could explain why some of the smallest islets appear to increase in intensity over the course of a glucose challenge.

Using a heated slide (raised on heated copper tubing) with spacers between the slide and coverslip helped to restrict slide motion and improved secretory responsiveness. Rapid-fire imaging maximized the chance to capture images free of motion blur throughout the animal breathing cycle, with a motorized stage used for tiling higher-resolution images of the pancreas. Moreover, we used high dynamic range imaging (12-bit, 4,096 levels of intensity) to capture without saturation the faintest as well as the brightest islets. In addition to maximizing the range of intensities of islets that could be captured, the increase of dynamic fluorescence range increased the number of usable imaged islets that could be captured without saturation of the signal.

Islet response *in vitro* has been extensively studied and islets are known to respond in a coordinated manner, secreting only a small fraction of their insulin content (36). The platform described in this report allows for the first time the ability to follow differences in the responses of individual islets *in vivo* in anesthetized mice. Notably, while some islets respond acutely, others seem unaffected by the glucose challenge. Islet distribution is similar in mice and humans (37-41), scattered throughout the head, tail and body of the pancreas. Using crude immunological assessments, it had been hypothesized that some islets might respond better than others based on their location within the pancreas itself (35). However, our data does not support this hypothesis since islets in very close proximity to one another were found to display dramatically different responses (see Figure 7B). Moreover, the close proximity of islets exhibiting drastically different secretory responses would also seem to discount the possibility that mechanical stresses blunt the response across an entire region of the pancreas. Alternatively, it has been hypothesized that differences in response between islets could result from innervation, vascularization or that some islets are inherently stronger in their secretory response to glucose (35). Conceivably, the first islets to sense a rise in blood glucose could secrete larger amounts of insulin, thereby inhibiting responses of islets further downstream. However, this hypothesis remains untested. Further, if innervation controls the magnitude of the response to glucose, denervating the pancreas should theoretically restore homogeneity to responses between islets.

Described here is a platform that allows for the testing of various hypotheses about what contributes to heterogeneity between islet insulin secretion *in vivo*. Moreover, islets can be compared in the normal state, or under pathological conditions, or under the influence of various pharmacological agents, nutrients, infused antibodies, or surgical intervention. The CpepSfGFP mice offer the opportunity to cross with other loss or gain of function genetic models in order to test specific molecular hypotheses. The platform described here constitutes a robust and practical integrative physiology model that can be used to compare insulin secretion at the level of the islet in the physiological and pathophysiological *in vivo* milieu.

## Acknowledgements

This study was supported by the Michigan Diabetes Research Center (NIH P30 DK020572, including the Animal Studies Cores and islet isolation laboratory) as well as the Michigan Nutrition Obesity Research Center (NIH DK089503) and the NIH (grants DK117821 to R.S. and DK048280 to P.A.). In addition, we would like to thank William McRoy (from the University of Michigan Undergraduate Research Opportunity Program), Dennis Larkin, and Kelli Rule for their assistance with the experiments as well as the Sandoval and Seeley Labs for their helpful discussions.

HFS contributed to the design of the experiment, carried out the experiments, analyzed the data co-wrote the manuscript, PA contributed to the design of the experiments, RJS contributed to the design of the experiments, their interpretation and CCM contributed to the design of the experiment and the computational framwewok, carried out the experiments, analyzed the data, and co-wrote the manuscript. All authors provided critical feedback, discussed the results and reviewed the final manuscript.

## Figure legends

**Supplemental figure 1: Diagram of the relational database generated for the time course comparison**

Schematic representation of the relational database allowing the identification of the islets at different time points and the export of the corresponding integrated densities for each islet over time.

The database integrates comparisons right before (t0) and right after (t0p) glucose infusion as well as at every subsequent time point. Each object gets an identifier in FIJI. A table of manually tracked objects gets generated to match the objects from time points to time points and allows the tables of integrated densities at each time point to be put in relation in the database.

