## Supplementary figures and images for "*In vivo* imaging of individual islets across the mouse pancreas reveals a heterogeneous insulin secretion response to glucose"

### Supplemental Figure 1

Supplemental Figure 1:

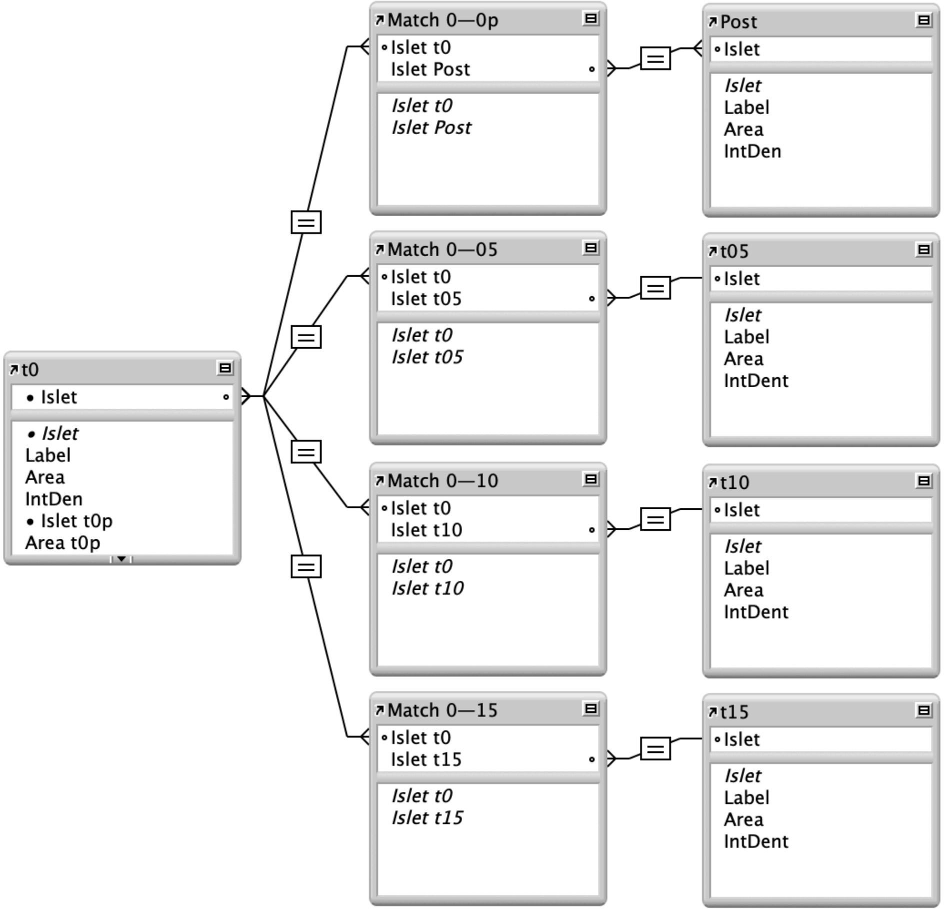
